# Determining the importance of the stringent response for methicillin-resistant *Staphylococcus aureus* virulence using a zebrafish model of infection

**DOI:** 10.1101/2023.07.12.548523

**Authors:** Naznin R. Choudhury, Lucy Urwin, Bartłomiej Salamaga, Lynne R. Prince, Stephen A. Renshaw, Rebecca M. Corrigan

## Abstract

*Staphylococcus aureus* is a bacterial pathogen that poses a major threat to human health. The ability of this bacterium to adapt to stresses encountered in the host is essential for disease. The stringent response is a signalling pathway utilised by all bacteria to alarm cells when stressed, and has been linked to the virulence of a number of species. This signalling pathway is controlled by the nucleotide alarmones guanosine tetra-(ppGpp) and pentaphosphate (pppGpp: collectively termed (p)ppGpp), produced in *S. aureus* by three synthetase enzymes: Rel, RelP and RelQ. Here, we used a triple (p)ppGpp synthetase mutant ((p)ppGpp^0^) to examine the importance of this signalling network for the survival and virulence of *S. aureus in vivo*. Using an established zebrafish larval infection model, we observed that infection with (p)ppGpp^0^ resulted in attenuated virulence, which was not due to a reduced ability of the mutant to replicate *in vivo*. Of the three (p)ppGpp synthetases, Rel was established as key during infection, but roles for RelP and RelQ were also observed. Zebrafish myeloid cell depletion restored the virulence of (p)ppGpp^0^ during systemic infection, indicating that (p)ppGpp is important for survival within host phagocytes. Primary macrophages infection studies, followed by *in vitro* tolerance assays to key innate immune effectors, demonstrated that (p)ppGpp^0^ was more susceptible to stressors found within the intracellular macrophage environment, with roles for all three synthetases implicated. Lastly, the absence of CodY, a transcription factor linked to the stringent response, significantly increased the tolerance of *S. aureus* to phagolysosomal-like stressors *in vitro*, but had no impact *in vivo*. Taken together, these results define the importance of the stringent response for *S. aureus* infection, revealing that (p)ppGpp produced by all three synthetases is required for bacterial survival within the host environment by mediating adaptation to the phagolysosome.

## Introduction

*Staphylococcus aureus* is a highly adaptable pathogen, with a large arsenal of virulence factors that allow it to colonise diverse sites within the human host. Upon infection, bacteria are subjected to harsh conditions due to changes in nutrient availability, pH and temperature, as well as the presence of an immune response. When faced with stresses, bacteria induce a conserved survival pathway termed the stringent response, which is coordinated by the nucleotide alarmones guanosine tetraphosphate (ppGpp) and guanosine pentaphosphate (pppGpp: collectively known as (p)ppGpp) [1]. (p)ppGpp is produced by the RelA/SpoT Homologue (RSH) protein family [2], with *S. aureus* encoding three of these synthetases: the long RSH enzyme Rel, which also has hydrolase activity, and the two monofunctional small alarmone synthetases (SAS) RelP and RelQ (Fig. 1A) [3]. Production of (p)ppGpp by these enzymes results in major changes within the bacterial cell, with alterations to numerous macromolecular activities such as transcription and translation [4–6]. These alterations lead to an inhibition of growth and a concurrent upregulation of stress adaptation and virulence factors, which ultimately allows a switch from active growth to a more stationary phenotype to aid bacterial survival. In *S. aureus*, the three (p)ppGpp synthetases respond to different environmental stresses to increase (p)ppGpp levels, with Rel sensing amino acid starvation via interactions with uncharged tRNA and the ribosome, while RelP and RelQ respond to cell wall and pH stress [4].

**Fig. 1.**
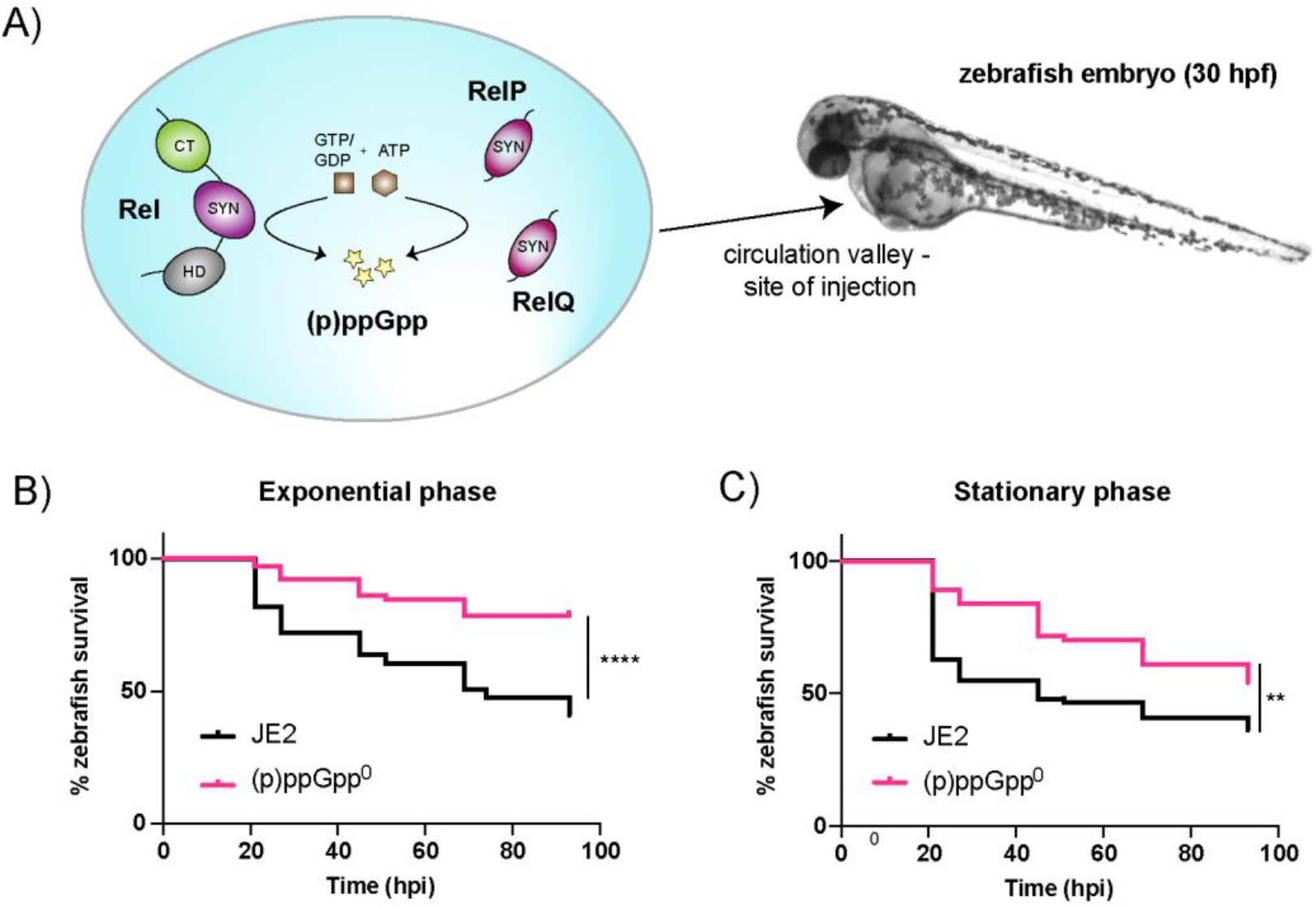
(p)ppGpp^0^ *S. aureus* strains have attenuated virulence in a systemic infection model. **A)** Schematic overview of the (p)ppGpp turnover enzymes in *S. aureus*. (p)ppGpp is produced by three enzymes, Rel, RelP and RelQ via the synthetase (SYN) domain. Rel is also capable of hydrolysing (p)ppGpp via the HD domain. Interactions between Rel and the ribosome occur via the C-terminal (CT) domain. *S. aureus* (blue circle) is injected into the yolk sac circulation valley of zebrafish embryos at 30 hpf. **B & C)** Survival of zebrafish larvae injected with *S. aureus* JE2 (black) and JE2 (p)ppGpp^0^ (pink) grown to **(B)** exponential and **(C)** stationary phase. Doses of 3000-4000 CFU of each strain were injected into the yolk sac circulation valley at 30 hpf to initiate a bloodstream infection. Survival was monitored until 93 hpi when the larvae reached 5.2 dpf. Pairwise comparisons (log-rank (Mantel-Cox) test) were as follows: B) (p)ppGpp^0^ versus wildtype, **** *P* < 0.0001, C) (p)ppGpp^0^ versus wildtype, ** *P* = 0.0048. Experiments were performed in quadruplicate for **(B)** and in triplicate **(C)**.

Activation of the stringent response is reported to contribute to the pathogenicity of a number of bacterial species. For example, a *Salmonella enterica* subspecies Typhimurium (p)ppGpp-null ((p)ppGpp^0^) mutant was unable to replicate in the mouse spleen after five days [7]. Similarly, the numbers of a *Mycobacterium tuberculosis rel* mutant recovered from murine lung and spleen tissues was 2-log lower than the wildtype over a 38-week period, implicating a requirement for Rel for long-term viability and chronic infection [8]. The absence of (p)ppGpp also affected the ability of *Enterococcus faecalis* to form biofilms in murine models of catheter-associated urinary tract infections [9]. For *S. aureus*, a methicillin-resistant *S. aureus* (MRSA) *rel* mutant formed cutaneous abscess lesions in mice that were over 13-times smaller than those formed by the wildtype [10]. Rel was also required for maintaining methicillin-sensitive *S. aureus* (MSSA) load in murine renal abscesses and for reducing mouse body weight [11]. This loss of body weight was dependent on the transcription factor CodY, which derepresses amino acid and virulence genes upon induction of the stringent response [12]. Together, these reports indicate that the stringent response is important for the virulence of a number of bacterial pathogens.

Zebrafish (*Danio rerio*) are a well-established animal model for various types of disease and infection. Despite being non-mammalian, zebrafish are jawed vertebrates with high genetic homology to humans - more than 80% of disease-associated genes have a human counterpart [13]. Furthermore, advantages over mammalian models include its rapid embryonic development, genetic manipulability and its transparency at the embryonic and larval stages, which allows for live imaging of developing zebrafish [14]. With a functional innate immune system by 30 hours post fertilisation (hpf) [15, 16], zebrafish larvae are also useful for studying host-pathogen interactions, as demonstrated by the numerous infection models that exist [17]. For example, we have previously developed systemic infection models to study the pathogenicity of *S. aureus* and *E. faecalis* within zebrafish [18, 19]. Studies such as these highlight how the zebrafish model can be used to further our knowledge of host-pathogen interactions and disease pathogenesis.

While reports indicate that the Rel synthetase is important for the survival of *S. aureus* in polymorphonuclear leukocytes (PMNs) [20], as well as in cutaneous abscess lesions and murine renal abscess models [10, 11], the importance of the entire signalling system, and the contribution of the two SAS enzymes, RelP and RelQ, to virulence is much less well understood. Here, we sought to use the versatility of the zebrafish model to establish the importance of (p)ppGpp for *S. aureus* systemic infection, extending previous findings relating to bacterial load in host organs to examine the contribution of the stringent response to host killing. Using the zebrafish model, we have determined that all three synthetases contribute to the virulence of *S. aureus*. The attenuated phenotype of a (p)ppGpp^0^ mutant was determined to be at least partly myeloid cell-dependent, as morpholino-mediated ablation of myeloid cells restored virulence. Moreover, we show a requirement for (p)ppGpp for survival of *S. aureus* within primary macrophages, with *in vitro* studies highlighting its importance for tolerance of phagolysosomal stressors. While the overproduction of (p)ppGpp has been suggested to support chronic and recurrent infections, here we observed that increased (p)ppGpp production led to higher tolerance of *S. aureus* to stressors *in vitro* but actually reduced bacterial virulence. Finally, deletion of the CodY transcription factor restored the survival defect of the (p)ppGpp^0^ mutant *in vitro*, but did not restore virulence. Altogether, this work further defines the importance of (p)ppGpp, and each of the three synthetases, for *S. aureus* growth, systemic infection and host death.

## Materials and methods

### Bacterial strains and culture conditions

*Escherichia coli* strains were grown in Luria Bertani broth (LB). *S. aureus* strains were grown in tryptic soy broth (TSB) at 37°C with aeration. Strains used in this study are listed in S1 Table and primers used are listed in S2 Table. Plasmids pALC2073-*relQ,* pCL55iTETr862-*rel* and pCL55iTETr862-*relP* were constructed by amplifying the respective genes using the primers listed in S2 Table. The resulting PCR products were digested and cloned into either pALC2073 or pCL55iTETr862 that had been digested with the same enzymes. All plasmids were initially transformed into *E. coli* strain XL1-Blue and sequences of all inserts were verified by fluorescence automated sequencing by Eurofins. The *S. aureus* expression plasmids pALC2073 and pALC2073-*relQ* were first electroporated into RN4220 before isolation and electroporation into JE2. The integrative pCL55iTETr862 plasmids were electroporated into RN4220 before being phage transduced into JE2 using Φ85. Φ85 was also used to move the *codY*::Tn transposon mutation into JE2 and JE2 (p)ppGpp^0^.

### Zebrafish strains and husbandry

Up to 5 days post fertilization (dpf) zebrafish are not protected under the Animals (Scientific Procedures) Act 1986. However, all work was carried out according to the stipulations set out in Project License P1A4A7A5E. London wildtype (LWT) strains were used for all zebrafish experiments. Adult zebrafish were maintained by staff at the University of Sheffield Bateson Centre Zebrafish Facility according to established standards [21]. Adult fish were kept at 28°C in a 14 hr/10 hr light/dark regime. Embryos/larvae were incubated at 28°C in E3 medium (0.5 mM NaCl, 17 µM KCl, 33 µM CaCl2, 33 µM MgSO4, 0.00005% methylene blue).

### Zebrafish embryo microinjections

At approximately 30 hpf, LWT zebrafish embryos were dechorionated and anaesthetised by immersion in 0.02% w/v buffered tricaine. The embryos were embedded in 3% w/v methylcellulose on a glass slide. 1 nl of bacterial suspensions were injected into the yolk sac circulation valley of the ≥ 30 embryos per condition using a pneumatic micropump (World Precision Instruments PV820), a micromanipulator (WPI) and a dissecting microscope. Following injection, embryos were recovered in fresh E3 and placed into individual wells of a 96-well plate. After 2 dpf, embryos are referred to as larvae. The larvae were monitored twice a day up to 93 hours post infection (hpi) and the number of dead larvae at each timepoint recorded. To confirm bacterial numbers in each injection, the same volume was ejected into 1 ml of PBS and the viable counts determined on tryptic soy agar (TSA) plates. Survival curves were generated using GraphPad Prism.

### Measurement of S. aureus growth in zebrafish

*S. aureus* cultures were injected into the yolk sac circulation valley of zebrafish embryos at 30 hpf. At each timepoint until 5.2 dpf, five live larvae and any dead larvae, as well as 200 µl of E3 medium were transferred to 0.5 ml microcentrifuge tubes containing 1.4 mm ceramic beads. Each larva was homogenised using a FastPrep-24™ 5G Homogeniser and homogenates were serially diluted and plated to determine bacterial load.

### Microinjection of morpholino-modified antisense oligonucleotides

One pmol of a morpholino-modified antisense oligonucleotide against the Pu.1 transcription factor [22] was injected into the yolk of one-cell stage zebrafish embryos, which were subsequently incubated at 28°C until injection with *S. aureus*. *S. aureus* cultures were injected into zebrafish embryos at 30 hpf. Larvae were maintained at 28°C, monitored twice a day up to 93 hpi (5.2 dpf) and the number of dead larvae at each timepoint recorded.

### Preparation of frozen stocks for macrophage infection

Bacterial strains were cultured overnight in TSB and diluted to an OD_600_ of 0.05. Diluted cultures were grown until mid-stationary phase (approx. 9 hrs), supplemented with 15% glycerol and stored at −80°C. Prior to infection, aliquots were thawed on ice, washed once with PBS and resuspended in RPMI 1640 containing no additional supplements. CFU/ml values of frozen stocks were routinely confirmed by plating and multiplicity of infection (MoI) values adjusted accordingly.

### Isolation and culture of human monocyte-derived macrophages (MDMs)

Leukocyte cones (supplied by NHS Blood and Transplant Service (NHSBT, UK) as anonymized samples from consenting donors) were used to isolate peripheral blood mononuclear cells (PBMCs) from human blood (day 0). Briefly, whole blood was separated by density centrifugation using Ficoll Paque Plus and the buffy layer (containing PBMCs) was extracted for further processing. Platelets were removed by low-speed centrifugation and Ammonium-Chloride-Potassium (ACK – Thermo Fisher) lysis buffer was used to lyse red blood cells. Isolated PBMCs were resuspended in RPMI 1640 medium containing 10% new-born calf serum, 1% L-Glutamine and 1% antibiotic-antimycotic solution. A cell count was performed and PBMCs were seeded into tissue culture plates at 2 x10^6^ cells/ml - this seeding density is estimated to provide 2 x 10^5^ cells/ml MDMs. After 24-48 hrs, the seeding medium was removed and replaced with RPMI 1640 containing 10% foetal bovine serum, 1% L-Glutamine and 1% antibiotic-antimycotic solution. This media was replaced every 3-4 days to promote the differentiation of MDMs. MDMs were used in experiments between 12-and 14-days post isolation and the supplemented RPMI 1640 media was replaced with RPMI 1640 that did not contain antibiotic-antimycotic solution at least 24 hrs prior to infection.

### Macrophage infection assays

PBMCs were seeded into 6-well tissue culture plates at 2 x10^6^ cells/ml. On day 12, MDMs were washed once with Hanks balanced salt solution (HBSS) and cells were dissociated by 20 min incubation with accutase at 37°C, followed by gentle cell scraping. Dissociated cells were pooled, centrifuged at 400 *x g* for 5 mins and resuspended in RPMI 1640 medium for cell counting. MDMs were seeded into 12-well tissue culture plates at 2 x10^5^ cells/ml and returned to tissue culture incubators prior to infection. On days 13-14, MDMs were washed once with HBSS and infected using frozen bacterial stocks at MoI 10. Plates were centrifuged at 277 *x g* for 2 min to synchronise infection and then incubated for 30 min at 37°C. After 30 min, cells were washed twice with ice-cold PBS to remove non-adherent bacteria and halt bacterial internalisation. Gentamicin was prepared in RPMI 1640 containing no additional supplements at 100 µg/ml and added to infected MDMs for 30 min at 37°C to kill extracellular bacteria. To measure bacterial killing, high-dose gentamicin (100 µg/ml) was replaced with RPMI 1640 containing 4 µg/ml gentamicin and 0.8 µg/ml lysostaphin. Infected MDMs were incubated at 37°C until 6 hpi and low-dose gentamicin/lysostaphin was removed. Cells were washed twice with PBS and intracellular bacteria enumerated.

### Tolerance assays

*S. aureus* overnight cultures were diluted to OD_600_ of 0.05 and grown to mid-exponential at 37°C with aeration at 200 rpm with antibiotics if required, including 50 ng/ml anhydrotetracycline (Atet) for the iTET-inducible strains. Once the desired optical density (approximately OD_600_ 0.35) was reached, the cultures were centrifuged at 4000 x *g* for 10 min and washed twice in sterile PBS. Antimicrobial compounds (20 mM itaconic acid, 100 mM H_2_O_2_ (from a 30% w/w stock) or 32 μM sodium hypochlorite/HOCl (from a stock containing 10-15% available chlorine) were added to OD_600_ 0.35 cultures (including antibiotics and Atet if required) and incubated at 37°C with aeration at 200 rpm. CFU were determinated at 0.5 or 1 hr after addition of each antimicrobial compound. Experiments were repeated up to ten times due to variation in survival between biological replicates.

### Statistics

Statistical analyses were performed using GraphPad Prism 9.0 software. Statistical differences between zebrafish larval survival experiments were evaluated using the Kaplan– Meier method and pairwise comparisons between survival curves were made using the log-rank (Mantel-Cox) test. For tolerance assays the normality of data sets were determined using the Shapiro-Wilk test. Differences in tolerance were then assessed using either Mann-Whitney test, or one-way ANOVA followed by Tukey’s multiple comparisons test or Kruskal-Wallis multiple comparison test, as indicated in the figure legends. Macrophage assays were analysed by one-way ANOVA with Dunnett’s multiple comparisons test.

## Results

### (p)ppGpp is important for S. aureus virulence in a zebrafish model

To determine the requirement of a functional stringent response for *S. aureus* virulence, zebrafish embryos were infected with either the community-acquired MRSA strain JE2, or a JE2 (p)ppGpp^0^ mutant [23]. Bacteria were injected into the bloodstream via the yolk sac circulation valley (Fig. 1A). Bacteria injected here enter the heart before more widespread dissemination throughout the bloodstream, culminating in bacteraemia [18]. A dose of approximately 3000 - 4000 CFU of wildtype JE2 led to 50% zebrafish mortality (Fig. 1). In contrast, the (p)ppGpp^0^ mutant killed significantly fewer larvae, which occurred with both exponentially-grown (Fig. 1B: *P* < 0.0001) or stationary phase (Fig. 1C: *P* = 0.0048) bacterial cultures. This approach confirms the usefulness of zebrafish larvae for studying *S. aureus* infection dynamics and establishes a role for (p)ppGpp in the virulence of *S. aureus*.

The JE2 (p)ppGpp^0^ strain grows similarly to wildtype under non-stressed conditions *in vitro* [23], however the mutant may have a replication defect *in vivo* explaining its decreased ability to cause death. To examine this, the *in vivo* bacterial growth kinetics for both the wildtype and the (p)ppGpp^0^ strain were elucidated by enumerating the bacterial CFUs over the course of the infection. Both strains were injected into zebrafish embryos and at each timepoint five live larvae, and any dead larvae, were homogenised and the CFU/larva determined (Fig. 2A, 2B). At 21 hpi, bacterial loads had increased from the initial inoculum of 10^3^ to between 10^5^ - 10^7^ for both JE2 and the (p)ppGpp^0^ mutant. This demonstrates that both were able to replicate within the larvae, although there were more dead larvae in the JE2-infected population. The (p)ppGpp^0^ mutant was isolated from larvae at numbers higher than 10^3^ from timepoints up to 69 hpi, suggesting that the (p)ppGpp^0^ mutant is also able to replicate later on during infection (Fig. 2B). Altogether, this suggests that while the (p)ppGpp^0^ mutant strain has attenuated virulence *in vivo*, it is still able to replicate in the host.

**Fig. 2.**
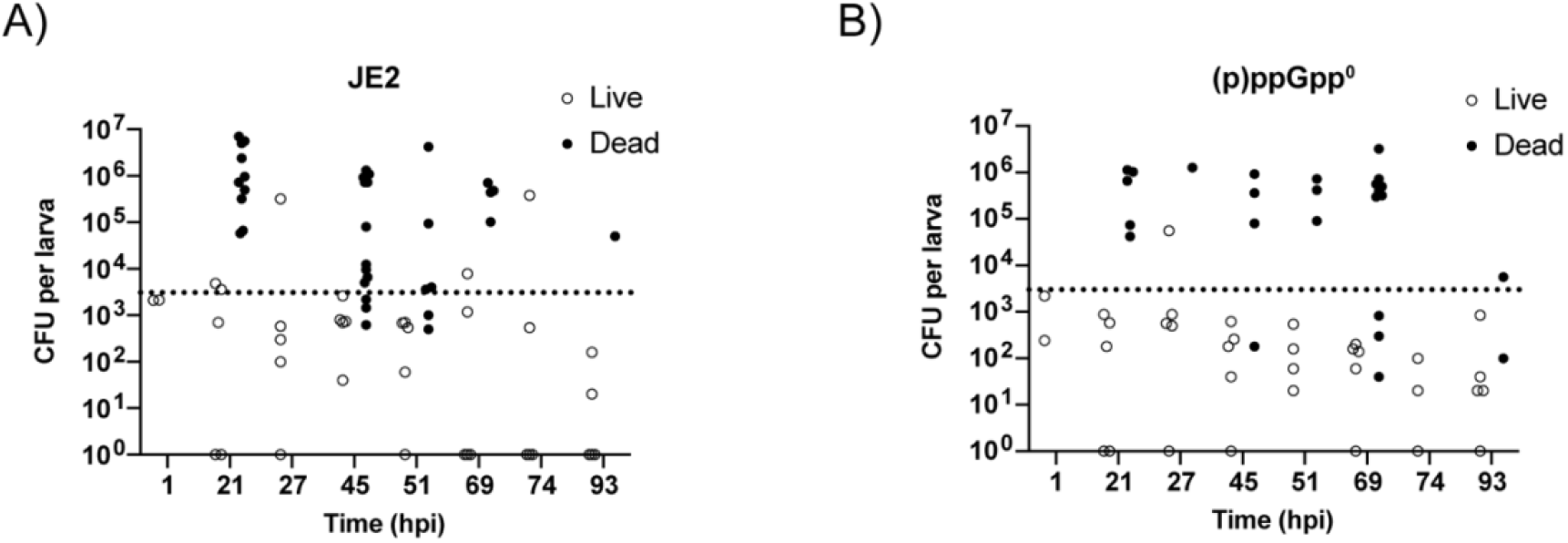
Wildtype and (p)ppGpp^0^ mutant both replicate *in vivo*. **A, B)** Growth of *S. aureus* JE2 **(A)** and (p)ppGpp^0^ **(B)** in zebrafish larvae after injection of 3000-4000 CFU (dotted line) into the bloodstream at 30 hpf. Five live larvae and any dead larvae were taken at the specified timepoints for CFU/embryo determination. Open circles represent live larvae and closed circles represent dead larvae. Survival was monitored until 93 hpi when the larvae reached 5.2 dpf. Experiment was performed in triplicate with one representative shown.

### Rel, RelP and RelQ all contribute to virulence of S. aureus

In *S. aureus*, (p)ppGpp is produced by the long bifunctional RSH enzyme Rel, as well as from the two SAS enzymes RelP and RelQ in response to different stresses [3]. To understand the contribution of the RSH versus the SAS enzymes to *S. aureus* infections, the virulence of JE2 and the (p)ppGpp^0^ mutant were first compared to JE2 Δ*relQP*, a strain with in-frame deletions of both SAS enzymes. Survival curves revealed that the Δ*relQP* mutant was able to kill larvae similarly to JE2 (Fig. 3A, 3E), suggesting that the presence of Rel is sufficient for virulence in this model. To confirm this, the (p)ppGpp^0^ mutant was complemented with full-length *rel* from the Atet-inducible integrative vector pCL55iTETr862 (iTET). Expression of Rel restored killing of the larvae to wildtype levels (Fig. 3B, 3E), while complementation with the single SAS enzyme *relP* did not (Fig. 3C, 3E). This confirms the importance of the Rel synthetase *in vivo*, as has been reported previously [10, 11].

**Fig. 3.**
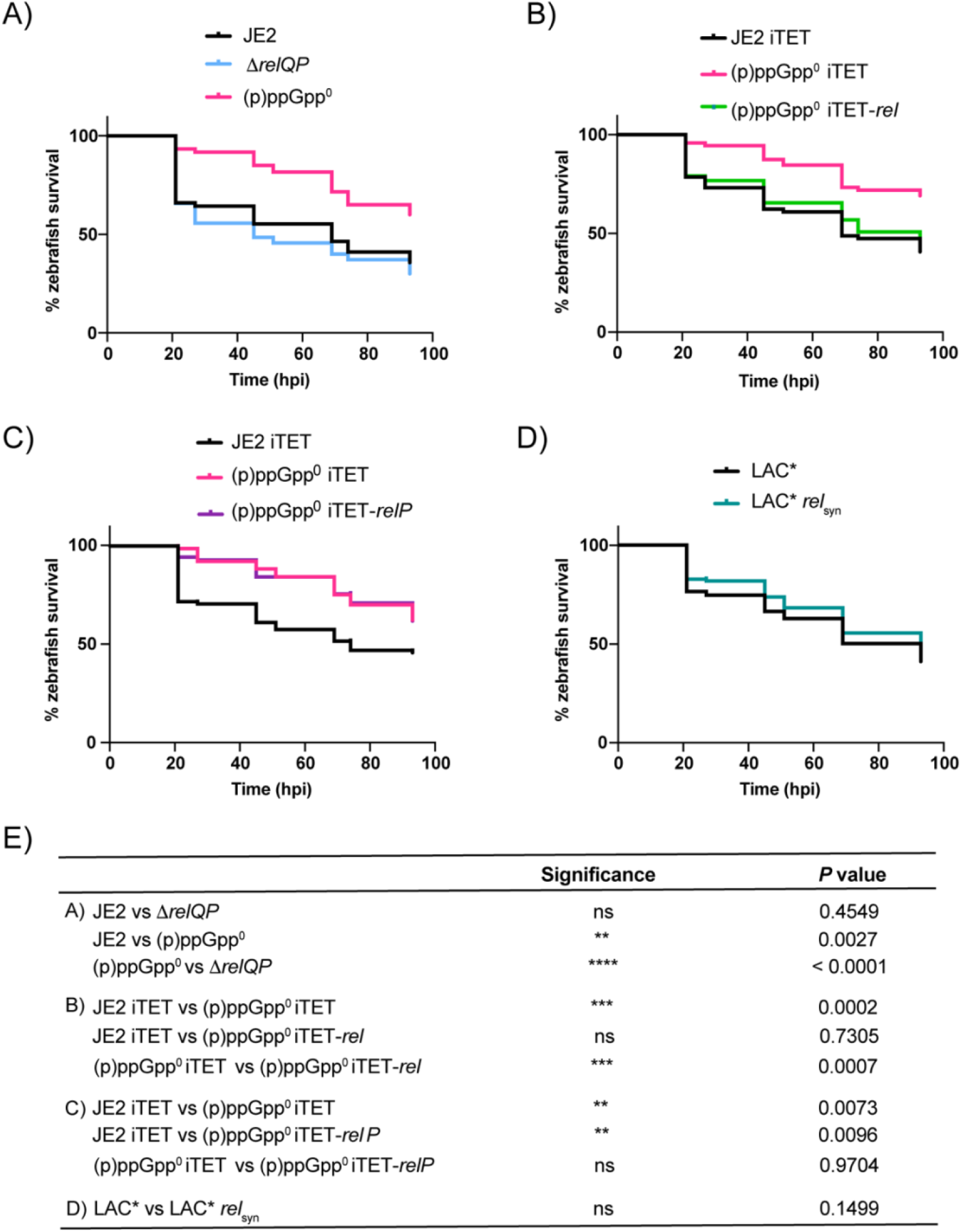
All three (p)ppGpp synthetases contribute to *S. aureus* virulence in a systemic zebrafish infection model. Survival of zebrafish larvae injected with *S. aureus* at 30 hpf. Survival was monitored until 93 hpi when the larvae reached 5.2 dpf. **A)** Injection of JE2 (black), (p)ppGpp^0^ (pink) and Δ*relQP* (blue) (dose 3000-4000 CFU). **B)** Injection of JE2 iTET (black), (p)ppGpp^0^ iTET (pink) and (p)ppGpp^0^ iTET-*rel* (green) (dose 3000-4000 CFU). **C)** Injection of JE2 iTET (black), (p)ppGpp^0^ iTET (pink) and (p)ppGpp^0^ iTET-*relP* (purple) (dose 3000-4000 CFU). **D)** Injection of LAC* (black) and LAC* *relsyn* (teal) (dose 1500 CFU). **E)** Statistical significance of A-D was determined by Log-rank (Mantel-Cox) test: ns, *P* > 0.05; \*\**P* < 0.01; *** *P* < 0.001; **** *P* < 0.0001. For A, B and C, experiments were performed in triplicate, while D was performed in quadruplicate.

While the above data indicate that the presence of Rel alone is sufficient for *S. aureus* virulence, we wanted to determine whether a strain containing the two SAS enzymes alone in the absence of Rel had a virulence defect. To establish this, we used a Rel mutant strain in which three conserved amino acids in the synthetase domain (Y308, Q309 and S310) are deleted, rendering it unable to produce (p)ppGpp. This leaves the hydrolase function intact, which is essential in strains encoding RelP and RelQ to prevent toxic accumulation of (p)ppGpp [11, 24]. This mutant was available in the LAC* background, a strain identical to JE2, except JE2 has been cured of the cryptic plasmid p01 [25]. A comparison of the virulence of this mutant to the wildtype LAC* revealed no difference in killing (Fig. 3D, 3E). This would suggest that while the presence of Rel alone is sufficient for infection with *S. aureus*, the combined level of (p)ppGpp produced by RelP and RelQ in LAC* *rel*syn is enough to compensate for the lack of the Rel synthetase activity. Altogether this indicates a role for all three enzymes, and not just Rel, in the virulence of *S. aureus*.

### The attenuated virulence of the (p)ppGpp^0^ mutant is myeloid cell-dependent

In the early stages of development, zebrafish larvae use myeloid cells to protect against infection [26]. To determine the contribution of myeloid cells to controlling the virulence of the JE2 (p)ppGpp^0^ mutant, both the wildtype and mutant strains were injected into embryos where myeloid cell depletion was induced. Here, a morpholino-modified antisense oligonucleotide was employed to transiently knockdown *pu.1*, encoding a transcription factor that is an integral component in the differentiation of pluripotent haematopoietic stem cells into cells of the myeloid lineage [27]. A knockdown of *pu.1* results in the delayed appearance of macrophages and neutrophils from 25 hpf to 48 hpf, and from 30 hpf to 36 hpf, respectively [15, 16, 28, 29]. At the one-cell stage of embryonic development, 1 pmol of the *pu.1* morpholino was injected into the yolk sac of the embryos, followed by injection of either JE2 or the (p)ppGpp^0^ mutant at 30 hpf into the circulation valley. Depletion of myeloid cells resulted in 100% killing of the larvae within 24 hpi and crucially, restored the virulence of the (p)ppGpp^0^ mutant to the same level as the wildtype (Fig. 4A). This indicates that myeloid cells are necessary for controlling *S. aureus* bloodstream infections, and that (p)ppGpp is required for the survival of *S. aureus* within these cells.

**Fig. 4.**
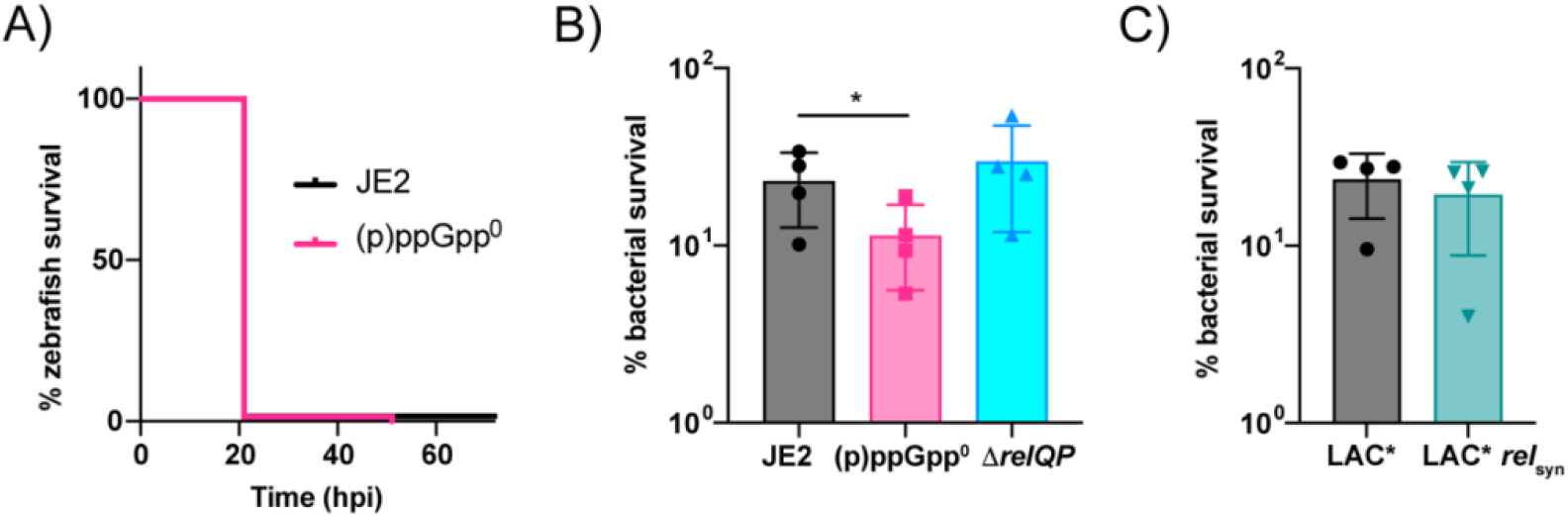
Macrophages are required to control *S. aureus* infection in a (p)ppGpp-dependent manner. **A)** Survival of Pu.1 knockdown zebrafish larvae injected with JE2 (black) and (p)ppGpp^0^ (pink) at doses of 3000-4000 CFU at 30 hpf into the circulation valley. 1 pmol of the Pu.1 morpholino was injected into the yolk of one-cell stage embryos. Survival was monitored until 93 hpi when the larvae reached 5.2 dpf. The experiment was performed in triplicate. Statistical significance was determined by Log-rank (Mantel-Cox) test: ns, *P* = 0.5275. **B, C)** Bacterial survival within human MDMs at 6 hpi. MDMs were infected with bacteria at MoI 10 for 30 min, before addition of 100 µg/ml gentamicin for 30 min to kill extracellular bacteria. Infected MDMs were lysed at 1 and 6 hpi and intracellular CFUs were used to calculate percentage bacterial survival at 6 h. Two technical repeats were performed using MDMs from each donor, with a total of 4 donors. The statistical significance of B was determined by one-way ANOVA with Dunnett’s multiple comparisons test: * *P* < 0.05.

### (p)ppGpp is required for the survival of S. aureus within primary macrophages

Previous work has demonstrated the importance of the (p)ppGpp synthetase Rel for survival of *S. aureus* within PMNs [20]. With our work showing that all three synthetases contribute to the virulence of *S. aureus in vivo*, we wished to understand more about the contribution of each to the survival of *S. aureus* within professional phagocytes. To this end, we monitored the ability of the (p)ppGpp^0^ mutant to survive within macrophages. Following internalisation, infected macrophages were incubated at 37°C for a further 6 hrs before surviving numbers were determined. In comparison to the wildtype, the (p)ppGpp^0^ mutant was significantly less able to survive the intracellular environment within macrophages (Fig. 4B).

To understand more about the individual contribution of Rel versus the SAS enzymes to this phenotype, we monitored the survival of both the Δ*relQP* and the *rel*syn mutants. A strain lacking RelP and RelQ survived just as well as the wildtype, indicating that Rel alone is sufficient to promote bacterial survival (Fig. 4B). This complements the zebrafish data, where Rel alone is sufficient for virulence (Fig. 3). The LAC* *rel*syn mutant, where the synthetase domain of Rel is inactivated, had a lower, but not statistically significant, survival rate than the wildtype, and was not killed as efficiently as the (p)ppGpp^0^ mutant, suggesting that while Rel plays a key role in this niche, RelP and RelQ also aid survival in its absence (Fig. 4C). Altogether, these data show that the reduced virulence observed in zebrafish is likely due to the inability of stringent response mutants to survive within macrophages and highlights a role for both the long RSH enzyme Rel and the SAS enzymes RelP and RelQ for responding to stress signals and producing sufficient (p)ppGpp to allow survival within these cells.

### The (p)ppGpp^0^ mutant is more susceptible to stress conditions found within a macrophage

Upon infection, *S. aureus* cells are phagocytosed by macrophages and incorporated into a phagolysosome. Here they encounter various insults including, but not limited to, low pH, reactive oxygen/nitrogen (ROS/RNS) species and antimicrobial peptides [30]. The main contributor to low pH is the proton-pumping v-ATPase, present on the membrane of phagolysosomes, though metabolites such as itaconic acid also contribute to this. Itaconic acid is produced in phagocyte mitochondria by aconitate decarboxylase, an enzyme that converts aconitic acid, a by-product of the Krebs cycle, to itaconic acid [31].

Previously, a methicillin sensitive *S. aureus* (MSSA) (p)ppGpp^0^ mutant exhibited susceptibility to H_2_O_2_ [5], while an MRSA (p)ppGpp^0^ mutant displayed reduced tolerance to HOCl [32]. These studies suggest that the stringent response is important for surviving stresses within the phagolysosome. To confirm this, JE2 and the (p)ppGpp^0^ mutant were exposed to HOCl, H_2_O_2_ and itaconic acid, and tolerance quantified (Fig. 5A-C). In keeping with previous observations, the (p)ppGpp^0^ mutant was 1-2 log more susceptible to HOCl and H_2_O_2_, and additionally showed a 1-log decreased tolerance to itaconic acid. To examine roles for each synthetase in combatting external stressors, the (p)ppGpp^0^ mutant was complemented with either the RSH enzyme Rel or the SAS enzyme RelP. Expression of RelP, while alone was unable to restore virulence in zebrafish (Fig. 3C), was sufficient to restore tolerance to both ROS stress and pH stress *in vitro* (Fig. 5D, 5E), while expression of Rel conferred tolerance to ROS stress only (Fig. 5D). These data are in keeping with previous reports suggesting roles for SAS enzymes in responding to pH stress [3, 33], as well as work indicating that expression of Rel is sufficient to combat ROS stress [32]. They also support the idea that the different classes of synthetase produce (p)ppGpp in response to different environmental stresses.

**Fig. 5.**
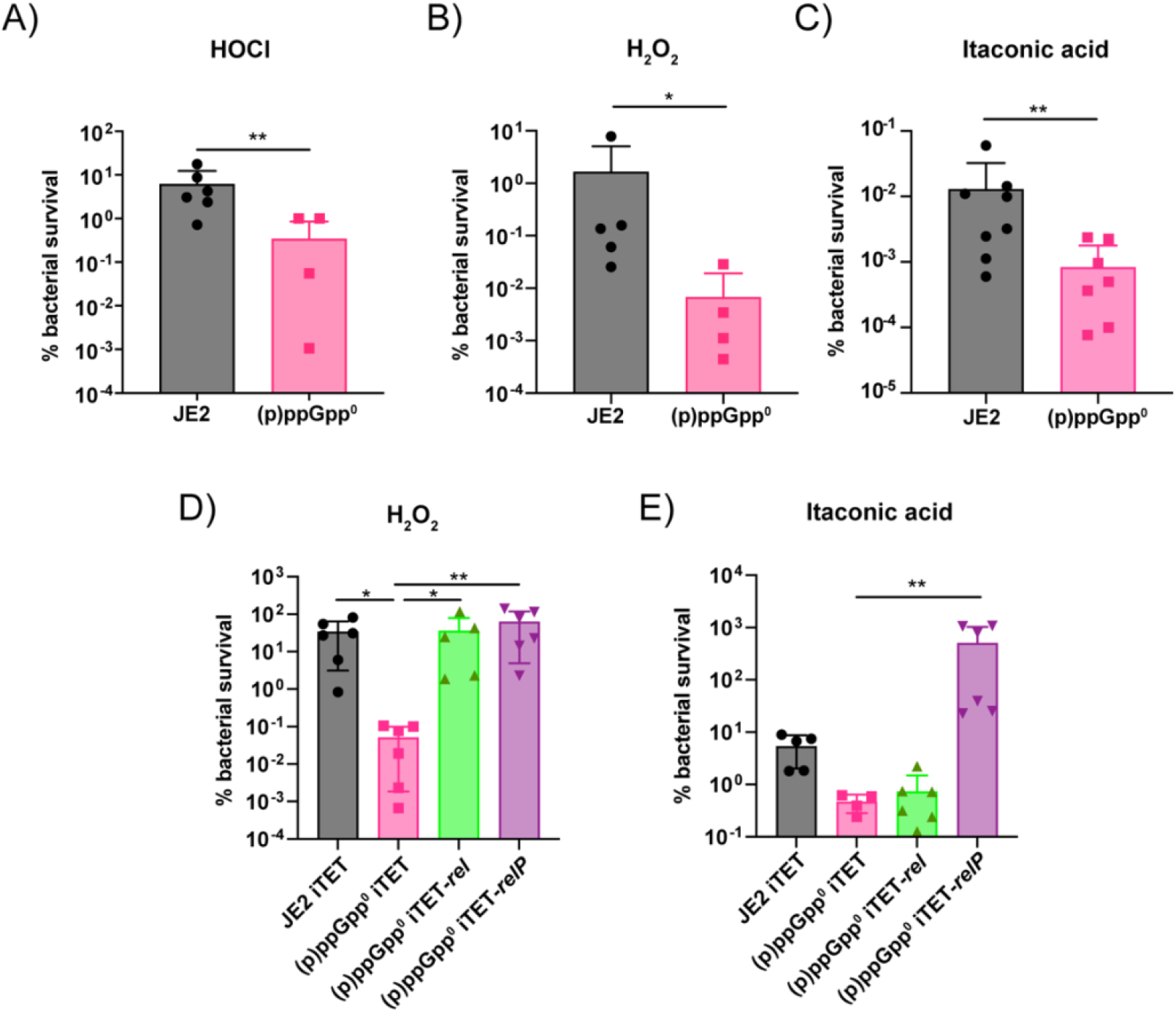
The (p)ppGpp^0^ mutant is less tolerant to stressors found within professional phagocytes. JE2 (black) and (p)ppGpp^0^ (pink) were grown to OD_600_ 0.35 in TSB. Cells were washed twice in PBS and exposed to **A)** 32 μM HOCl, **B)** 100 mM H_2_O_2_, or **C)** 20 mM Itaconic acid for 1 hr at 37°C before CFU determination. Percentage bacterial survival with mean and standard deviation are plotted. Statistical analysis performed using Mann-Whitney test. *\* P <* 0.05, ** *P <* 0.01. **D, E)** JE2 iTET (black), (p)ppGpp^0^ iTET (pink), (p)ppGpp^0^ iTET-*rel* (green) and (p)ppGpp^0^ iTET-*relP* (purple) were grown to an OD_600_ of 0.35 in the presence of 50 ng/ml Atet. Cultures were washed twice in PBS and exposed to **D)** 100 mM H_2_O_2_ or **E)** 20 mM itaconic acid for 30 min at 37°C and the CFU/ml was determined. Percentage bacterial survival with mean and standard deviation are plotted. Statistical analysis performed using a Kruskal-Wallis test followed by a Dunn’s multiple comparison test, *\* P <* 0.05, ** *P <* 0.01.

### Overproduction of (p)ppGpp confers tolerance to stress conditions in vitro but reduces virulence

The overproduction of (p)ppGpp in a clinical *S. aureus* strain has been associated with a persistent infection that did not respond well to antibiotic therapy [34, 35]. As (p)ppGpp acts to aid bacteria in surviving stresses, excess (p)ppGpp may serve to provide enhanced protection. Thus, we investigated how (p)ppGpp overproduction affects the survival of *S. aureus* in the presence of H_2_O_2_, HOCl and itaconic acid. Here, we introduced the Atet-inducible multi-copy plasmid pALC2073-*relQ* into JE2 and measured bacterial survival upon overproduction of (p)ppGpp. In the presence of all three stressors, excess (p)ppGpp led to an increase in survival in comparison to the JE2 empty vector strain (Fig. 6A-C). These results indicate that surplus (p)ppGpp has a protective effect *in vitro* and could contribute to the survival of *S. aureus* in stress conditions found within a macrophage.

**Fig. 6.**
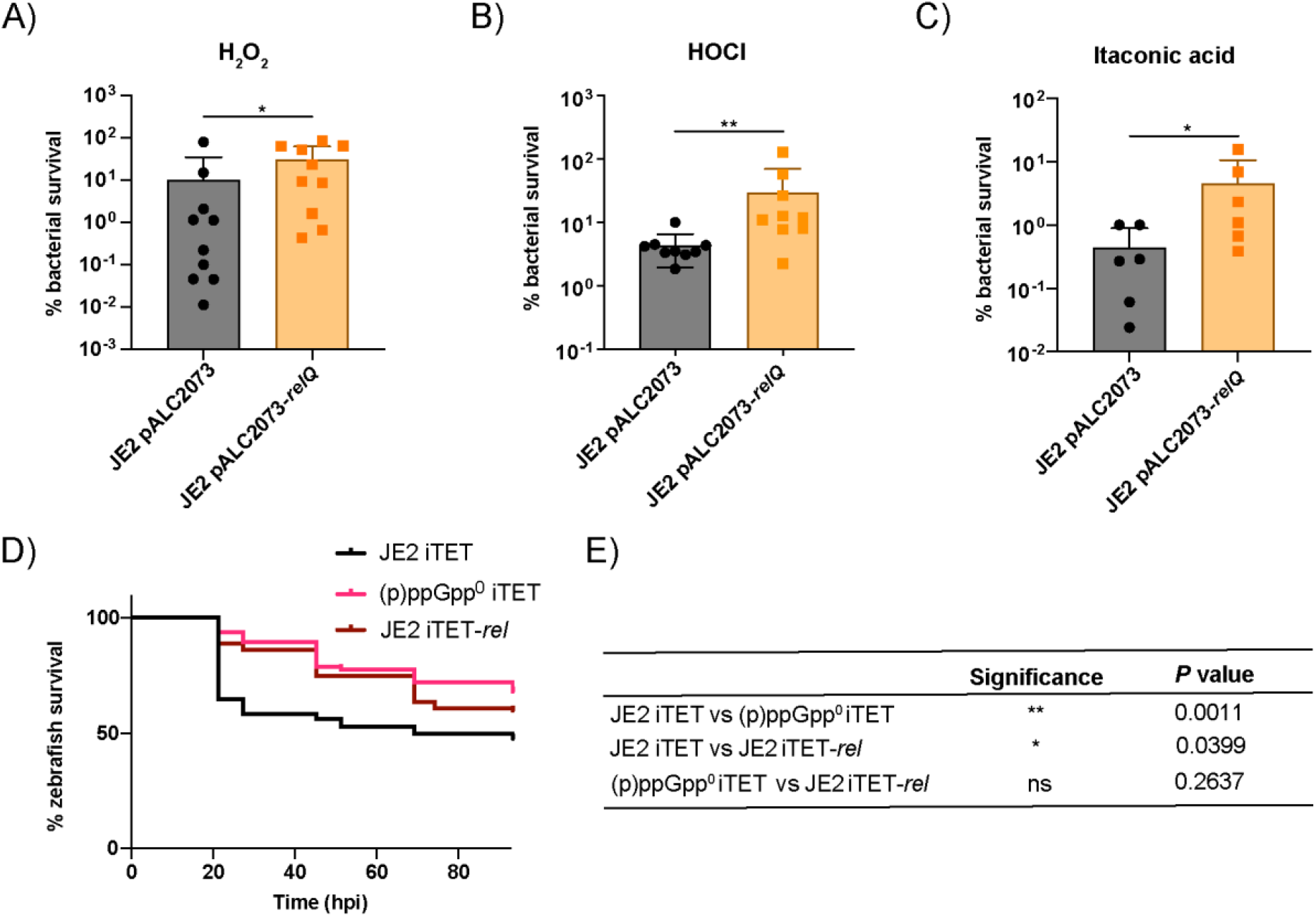
The tolerance of *S. aureus* to stressors found within professional phagocytes is increased in the presence of excess (p)ppGpp *in vitro* but virulence is reduced. **A - C)** JE2 pALC2073 (black) and JE2 pALC2073-*relQ* (orange) were grown to an OD_600_ of 0.35 in the presence of 50 ng/ml Atet. Cultures were washed twice in PBS and exposed to **A)** 100 mM H_2_O_2_, **B)** 32 μM HOCl or **C)** 20 mM itaconic acid for 30 min at 37°C after which the CFU/ml was determined. Percentage bacterial survival with mean and standard deviation are plotted. Statistical analysis performed using Mann-Whitney test. *\* P <* 0.05, ** *P <* 0.01. **D, E)** Survival of zebrafish larvae injected with *S. aureus*, JE2 iTET (black), (p)ppGpp^0^ iTET (pink), and the (p)ppGpp overproduction strain JE2 iTET-*rel* (brown) at doses of 3000-4000 CFU at 30 hpf into the circulation. Experiments were performed in triplicate. Statistical significance (E) was determined by Log-rank (Mantel-Cox) test: ns, *P* > 0.05; * *P* < 0.05, \*\**P* < 0.01.

Following this, we were curious to examine how overproduction of (p)ppGpp would impact the virulence of *S. aureus*. The (p)ppGpp overproduction strain JE2 iTET-*rel*, where a second copy of *rel* is integrated into the *S. aureus* genome, was injected into zebrafish embryos alongside the empty vector-containing controls. Here, overproduction of (p)ppGpp killed fewer larvae compared to the wildtype (Fig. 6D, 6E). While the *in vitro* data show that excess (p)ppGpp increases bacterial survival in macrophage-like conditions, *in vivo* assays suggest that (p)ppGpp overproduction does not improve virulence.

### Deletion of codY restores the survival defect of the (p)ppGpp^0^ mutant in vitro but does not impact virulence

Under nutrient-rich conditions, the CodY transcription factor represses genes related to nutrient acquisition and stress, including those involved in nitrogen and amino acid metabolism, as well as some virulence-associated genes [12]. In *S. aureus*, this repression requires GTP and branched-chain amino acids as CodY cofactors. During the stringent response, (p)ppGpp levels rise and cellular GTP levels fall. This is due to the consumption of GTP during the production of (p)ppGpp, in addition to the active inhibition of enzymes in the GTP synthesis pathway by (p)ppGpp [4]. This leads to the derepression of CodY and thus, the expression of stress-related genes in order to cope with the change in environment. As GTP levels in *S. aureus* are increased in strains lacking (p)ppGpp [23], we hypothesised that the continued repression of the CodY regulon by GTP-bound CodY in the (p)ppGpp^0^ mutant could be responsible for the decreased virulence phenotype observed in zebrafish larvae.

To investigate this, a *codY* mutant was introduced into both the wildtype and (p)ppGpp^0^ backgrounds, and the strains first exposed to itaconic acid and H_2_O_2_. In both the wildtype and the (p)ppGpp^0^ backgrounds, deleting *codY* rendered cells considerably more tolerant to stress (Fig. 7A, 7B), suggesting that the expression of genes previously repressed by CodY are beneficial for the survival of *S. aureus in vitro.* To examine the importance of CodY during systemic infection, zebrafish embryos were injected with the JE2 *codY*::Tn and (p)ppGpp^0^ *codY*::Tn mutants. In contrast to the *in vitro* results however, deleting *codY* did not rescue the attenuated virulence phenotype of the (p)ppGpp^0^ strain (Fig. 7C). This suggests that while inducing the CodY regulon is sufficient for enabling bacterial stress survival *in vitro*, it is not enough to restore virulence *in vivo* in a systemic infection model and suggests that processes regulated by (p)ppGpp independent of its connection with the CodY regulon are important for virulence of *S. aureus*.

**Fig. 7.**
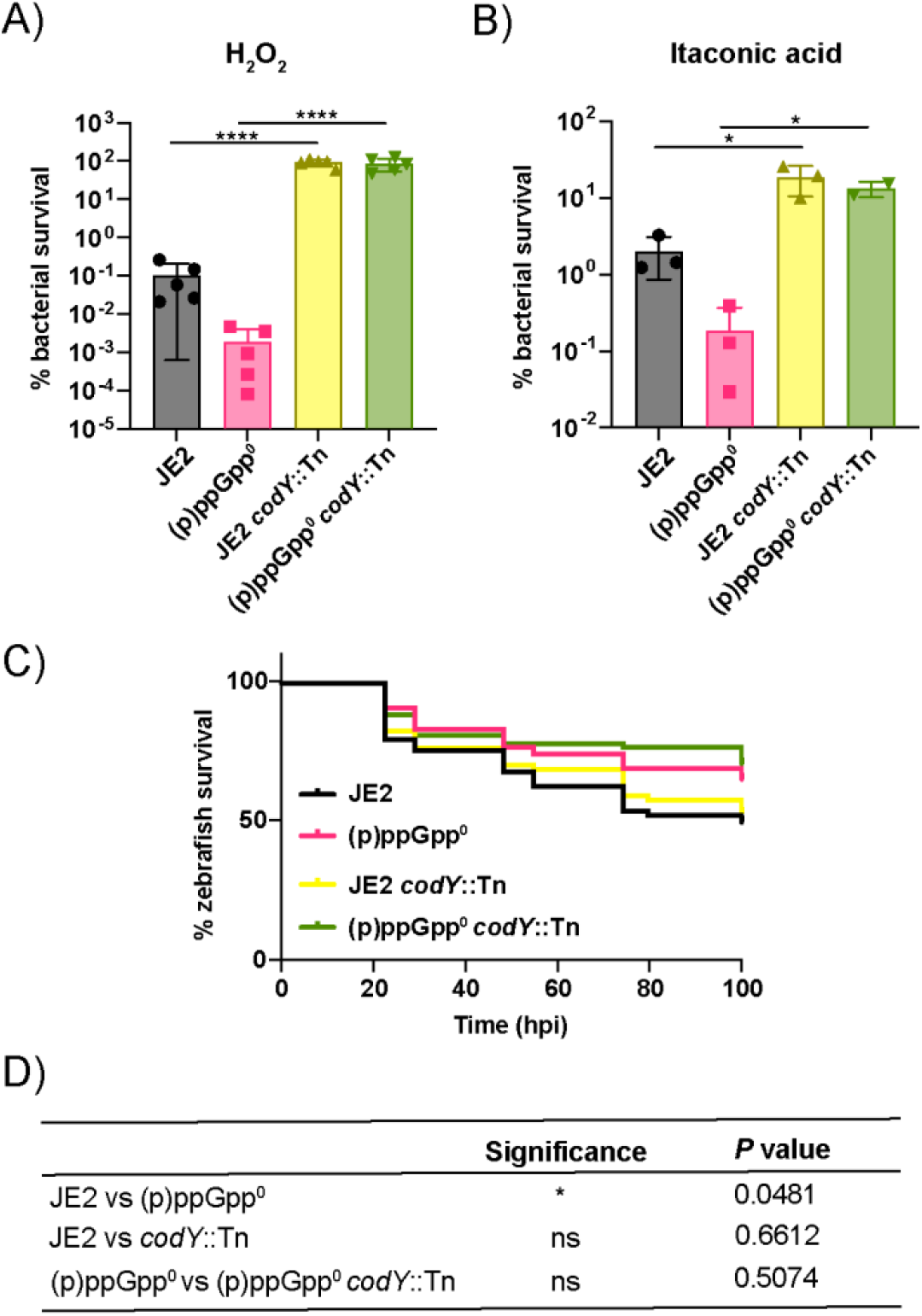
Deleting *codY* restores tolerance *in vitro* but does not affect virulence. **A, B)** Susceptibility of JE2 (black), (p)ppGpp^0^ (pink), JE2 *codY*::Tn (yellow) and (p)ppGpp^0^ *codY*::Tn (light green) to **A)** 100 mM H_2_O_2_ or **B)** 20 mM itaconic acid. Percentage bacterial survival with mean and standard deviation are plotted. Statistical analysis was performed using one-way ANOVA with Tukey’s multiple comparisons test. * *P* < 0.05, **** *P* < 0.0001. **C, D)** Survival of zebrafish larvae injected with *S. aureus*, comparing JE2 (black), (p)ppGpp^0^ (pink), JE2 *codY*::Tn (yellow) and (p)ppGpp^0^ *codY*::Tn (light green) at doses of 3000-4000 CFU at 30 hpf into the circulation. Survival was monitored until 93 hpi when the larvae reached 5.2 dpf. Statistical significance (D) was determined by Log-rank (Mantel-Cox) test: ns, *P* > 0.05; * *P* < 0.05. The experiment was performed in triplicate.

## Discussion

Upon infection environmental conditions become adverse, requiring stress responses such as the stringent response to modify cellular behaviour to maximise the chances of bacterial survival. Multiple studies have demonstrated the contribution of the staphylococcal stringent response to survival, including potential roles in: persistence [36, 37]; antibiotic resistance [35, 38, 39]; antibiotic tolerance [3]; immune evasion [20]; biofilm formation [40]; tolerance to oxidative stress [5, 32] and the development of murine pyelonephritis [11]. Thus, this study sought to systematically examine the importance of the stringent response, and each (p)ppGpp synthetase, for *S. aureus* virulence.

Our data shows that (p)ppGpp is required during systemic staphylococcal infection, as demonstrated by the attenuation of virulence for the (p)ppGpp^0^ strain (Fig. 1). During nutrient starvation, (p)ppGpp production and subsequent GTP depletion are required for the derepression of amino acid transport and synthesis genes via the transcription factor CodY [12]. This led us to hypothesise that the absence of (p)ppGpp could lead to a lack of nutrient acquisition and a subsequent growth defect *in vivo*. However, this was not the case, as both the wildtype and the (p)ppGpp^0^ mutant were able to replicate *in vivo* (Fig. 2). In agreement with this, cutaneous abscess formation by an *S. aureus rel* mutant was previously observed to be diminished, however the CFU per abscess was similar to the wildtype [10]. This is in contrast to the lower bacterial load of an MSSA *rel*_syn_ mutant recovered from murine kidneys [11], indicating that differences in bacterial load may occur at specific tissue sites.

The ability of the *S. aureus* (p)ppGpp^0^ mutant to replicate *in vivo* (Fig. 2), coupled with the observation that deleting the CodY repressor does not restore killing (Fig. 7), suggests that the absence of (p)ppGpp result in a survival or virulence defect, rather than a growth defect due to a lack of nutrient acquisition. This hypothesis is supported by multiple studies demonstrating an impact of the stringent response on virulence [9, 41–43]. By transiently depleting zebrafish embryos of myeloid cells, we demonstrated that the virulence of the (p)ppGpp^0^ mutant could be restored (Fig. 4A), supporting the idea that phagocytes are required for controlling *S. aureus* infection. As neutrophils are the most abundant circulating phagocyte [44], and thus are often the first immune cells to infiltrate a site of infection, further studies using this cell type are needed to examine the broader importance of the stringent response for infection.

The requirement of (p)ppGpp to survive ROS stress has been noted previously [32]. We extend this by showing that the stringent response is also necessary for tolerating itaconic acid, which may contribute to survival within macrophages. *In vivo*, itaconic acid has functions in addition to modulating the pH. It can have antimicrobial effects by inhibiting isocitrate lyase, a major component of the glyoxylate shunt that is an important pathway for optimal growth of bacteria [45–47]. Moreover, the charged conjugate base itaconate acts to reduce inflammation during *S. aureus* ocular infection by modulating NRF2/HO1 signalling and inhibiting the NLRP3 inflammasome [48]. Thus, future work is necessary to understand how the stringent response is required to modulate the anti-inflammatory roles of itaconic acid.

Previous transcriptome analysis has revealed that the *S. aureus* phenol-soluble modulin (PSM) cytotoxins are regulated via Rel independently of CodY [20], which, because of the role of PSMs in phagolysosomal escape [49], may contribute to the necessity of (p)ppGpp for survival in macrophages. Thus, it appears that the requirement for (p)ppGpp may be multi-factorial, where it could be needed for both surviving ROS and pH stress within the macrophage and for promoting escape via the PSMs. In the future it would be interesting to use microscopy to visualise the formation of the phagolysosome via recruitment of lysosome-associated membrane glycoproteins (LAMP) proteins. This would allow for visualisation of the bacteria within the phagolysosome and could be coupled with live-cell imaging to visualise phagolysosome escape.

This study demonstrates that all three (p)ppGpp synthetases contribute to the virulence of *S. aureus*. The importance of Rel, revealed by Geiger and colleagues [11, 20], is corroborated by our studies where we show that both the expression of *rel* in a (p)ppGpp^0^ mutant, and the presence of *rel* alone in a Δ*relQP* mutant, is sufficient to maintain wildtype levels of virulence (Fig. 3). This is then extended to show that both RelP and RelQ were sufficient for virulence (Fig. 3) and partial survival within macrophages in the absence of Rel (Fig. 4C). Due to the roles of RelP and RelQ in responding to cell wall and pH stress [4], and the fact that low pH is a condition encountered by pathogens following phagocytosis, it is not surprising that RelP and RelQ may play an important role in producing the (p)ppGpp required for the survival of *S. aureus in vivo*.

The impact of the stringent response on antimicrobial therapy outcomes in clinical settings has been demonstrated recently with the identification of MRSA and *Enterococcus* isolates from cases of persistent bacteraemia with constitutively active stringent responses [35, 50]. These two clinical isolates had reduced (p)ppGpp hydrolysis, reduced growth and increased antibiotic tolerance. Here, we observed that the overproduction of (p)ppGpp increased the tolerance of *S. aureus* to acid and ROS stress *in vitro* (Fig. 6A-C), however excess (p)ppGpp reduced *S. aureus* virulence (Fig. 6D). Likewise, Gao and colleagues found that a Rel hydrolase domain mutation led to a permanently activated stringent response in a clinical *S. aureus* strain isolated from a persistent infection. When the mutation was recapitulated in the laboratory, the strain also displayed attenuated virulence in a *Galleria mellonella* model [35]. This highlights the importance of regulating levels of (p)ppGpp within bacteria, and supports the idea that while (p)ppGpp overproduction may increase long-term persistence *in vivo* [35], it has a negative impact on virulence. Infection models lasting longer that 93 hpi would be required to delve into the role of (p)ppGpp in chronic infection further [36, 37].

Regulation by (p)ppGpp is interlinked with the transcription factor CodY. Previous studies have reported that introducing a *codY* deletion into *rel* mutant strains of both *S. aureus* and *Listeria monocytogenes* enhanced the ability of the strains to survive within phagocytes [11, 51]. Similarly, we revealed that both the JE2 *codY*::Tn and (p)ppGpp^0^ *codY*::Tn mutants were better able to withstand exposure to itaconic acid and H_2_O_2_ (Fig. 7A, 7B). Interestingly, this did not improve virulence (Fig. 7C). This is noteworthy, given that CodY regulates many virulence-associated genes, for example the virulence regulator *agr* [52, 53]. In *E. faecalis* and *Streptococcus agalactiae*, the absence of CodY resulted in reduced virulence in mice [9, 54]. However, when a *codY* deletion was introduced to a (p)ppGpp^0^ mutant, virulence was restored [9]. It is still unclear why some studies report hypervirulent Δ*codY* mutants while others report attenuated virulence. This is likely due to differences in species, strain and the infection models tested. As the (p)ppGpp and CodY regulatory networks are complex and intertwined, it may be necessary to investigate the expression of the CodY regulon during infection of zebrafish larvae to further understand this.

Altogether, this study has demonstrated that (p)ppGpp produced by all three synthetases contributes to *S. aureus* virulence, most likely through a need for surviving and escaping the harsh conditions within a phagolysosome. This work develops our understanding of the complexities of the (p)ppGpp regulatory pathways, which can inform the development of more effective stringent response inhibitors for the treatment of infections caused by strains such as MRSA.

## Acknowledgements

The work was supported by a Sir Henry Dale Fellowship jointly funded by the Wellcome Trust and the Royal Society (104110/Z/14/A to R.M.C.) and a Lister Institute Research Prize 2018 (to R.M.C.). The NHS Blood and Transplant have provided material in support of the research. This report is independent research. The views expressed in this publication are those of the author(s) and not necessarily those of NHS Blood and Transplant. The authors thank the Bateson Centre aquaria staff for their assistance with zebrafish husbandry. Josie Gibson, Joshua Sutton, Amy Tooke and Amy Lewis are acknowledged for help and advice on zebrafish work.

## SUPPLEMENTAL

**S1 Table.** Bacterial strains used in this study

| Strain | Relevant features | Reference |
| --- | --- | --- |
| *Escherichia coli* strains |  |  |
| XL1-Blue | Cloning strain: TetR | Stratagene |
| RMC0139 | pALC2073 in XL1-Blue: multi-copy vector: CarbR | [55] |
| RMC0546 | pALC2073-*relQ* in XL1-Blue: CarbR | This study |
| RMC0116 | pCL55iTETr862 (iTET) in XL1-Blue: single-copy, integrative vector: CarbR | [56] |
| RMC0468 | pCL55iTETr862-*rel* in XL1-Blue: CarbR | This study |
| RMC0469 | pCL55iTETr862-*relP* in XL1-Blue: CarbR | This study |
| *Staphylococcus aureus* strains |  |  |
| JE2 | CA-MRSA USA300 strain LAC derivative, lacking plasmids p01 and p03. Erm sensitive | [25] |
| RMC903 | JE2 $\Delta relQP$ JE2 with in-frame deletions in *relQ*, and *relP* | [23] |
| RMC905 | JE2 $\Delta relQPA$: JE2 with in-frame deletions in *relQ*, *relP* and *rel*: ((p)ppGpp<sup>0</sup>) | [23] |
| RMC1856 | JE2 pCL55iTETr862 (iTET): CamR | This study |
| RMC1858 | JE2 (p)ppGpp<sup>0</sup> iTET: CamR | This study |
| RMC1870 | JE2 (p)ppGpp<sup>0</sup> iTET-*rel*: CamR | This study |
| RMC1871 | JE2 (p)ppGpp<sup>0</sup> iTET-*relP*: CamR | This study |
| RMC1836 | JE2 pALC2073: CamR | This study |
| RMC1861 | JE2 pALC2073-*relQ*: CamR | This study |
| RMC2010 | JE2 iTET-*rel*: CamR | This study |
| LAC* | CA-MRSA USA300 strain LAC derivative, lacking<br>plasmid p03. Erm sensitive | [57] |
| NE1555 | JE2 *codY*::Tn. Strain with transposon insertion in *codY*:<br>ErmR | [25] |
| RMC1783 | JE2 *codY*::Tn – transduced into fresh JE2: ErmR | This study |
| RMC2014 | JE2 (p)ppGpp<sup>0</sup> *codY*::Tn: ErmR | This study |

Antibiotics were used at the following concentrations - for *E. coli* cultures: carbenicillin (CarbR) 50-150 µg/ml. IPTG was used at 1 mM. For *S. aureus* cultures: chloramphenicol (CamR) 7.5 or 10 µg/ml for pCL55iTETr862 or pALC2073 respectively; erythromycin (ErmR) 10 µg/ml; Atet 50 ng/ml.

**S2 Table.**
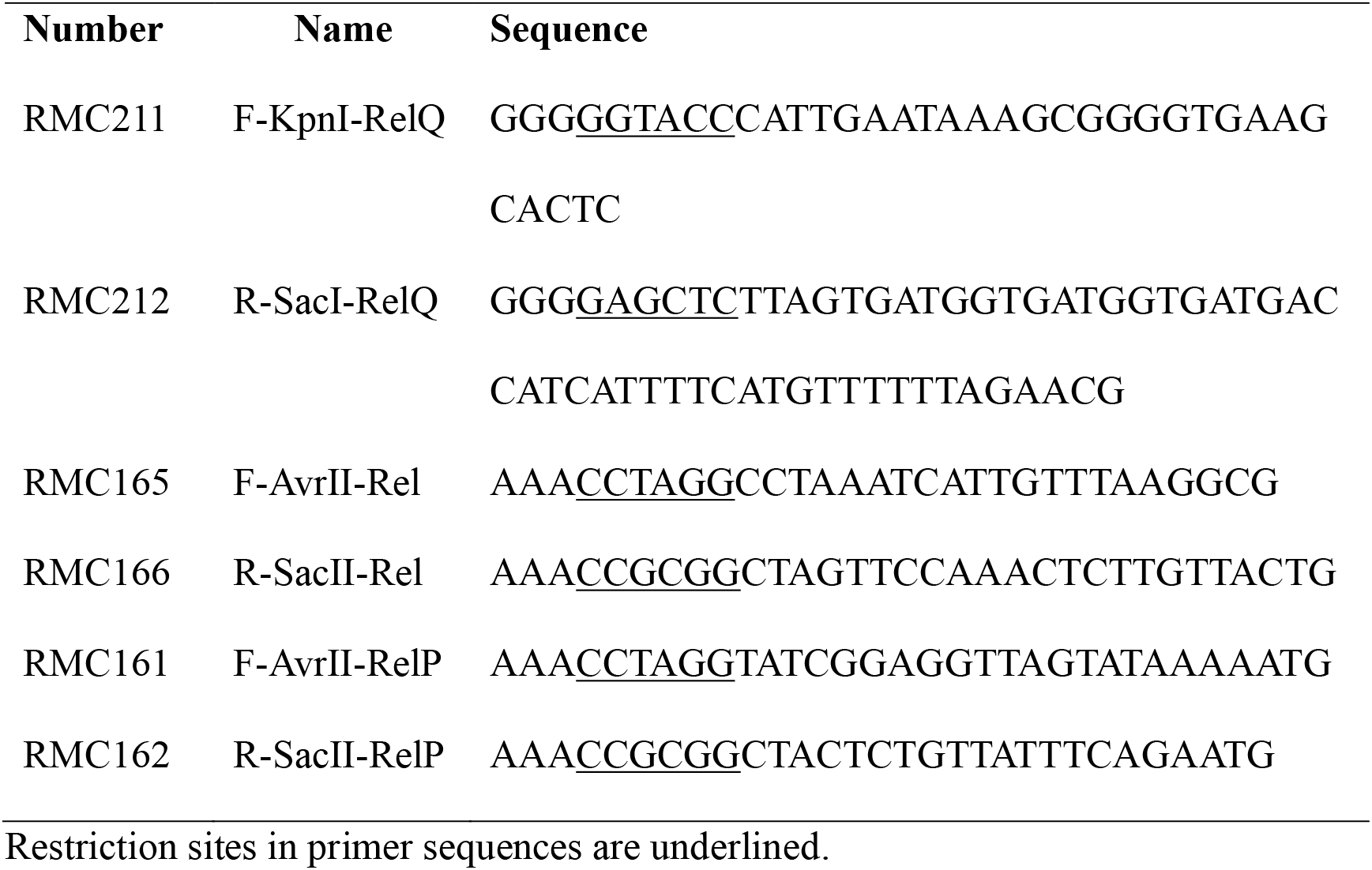
Primers used in this study

## Notes

### Competing Interest Statement

The authors have declared no competing interest.

